# Surgical procedure for implantation of Opto-Array in nonhuman primates

**DOI:** 10.1101/2022.10.04.510884

**Authors:** Reza Azadi, Simon Bohn, Mark A. G. Eldridge, Arash Afraz

## Abstract

Optogenetics allows precise temporal control of neuronal activity in the brain. Engineered viral vectors are routinely used to transduce neurons with light-sensitive opsins. However, reliable virus injection and light delivery in animals with large brains, such as nonhuman primates, has proven challenging. The Opto-Array is a novel yet simple device that is used to deliver light to extended regions of surface cortex for high-throughput behavioral optogenetics in large brains. Here we present protocols for surgical delivery of virus and implantation of the Opto-Array in two separate surgeries in a rhesus monkey’s inferior temporal cortex. As a proof of concept, we measured the behavioral performance of an animal detecting cortical optogenetic stimulations with different illumination power and duration using the Opto-Array. The animal was able to detect the optogenetic stimulation for all tested illumination powers and durations. A regression analysis also showed both power and duration of illumination significantly modulate the detectability of the optogenetic stimulation. The outcome of this approach is superior to the standard practice of injecting and recording through a chamber for large areas of surface cortex. Moreover, the chronic nature of the Opto-Array allows perturbation of neuronal activity of the same site across multiple sessions because it is highly stable, thus data can be pooled over months. The detailed surgical method presented here makes it possible to use optogenetics to modulate neuronal activity across large regions of surface cortex in the nonhuman primate brain.

## 1. Introduction

Optogenetic methods have revolutionized systems neuroscience (Tremblay et al., 2020). Optogenetics allows modulation of neuronal activity with precise spatial and temporal resolution, in a cell-type specific and/or projection specific manner. We recently developed the Opto-Array, a chronically implantable array of 24 LEDs and a thermal sensor configured in a 5 × 5 matrix for delivering light to large brains, specifically nonhuman primates. Each LED is able to illuminate ∼1 mm^2^ of the cortical surface allowing illumination of up to ∼5 × 5 mm of cortex (Rajalingham et al., 2021). The chronic nature of the Opto-Array allows reliable perturbation of neural activity at the same site in electrophysiology and behavioral studies, in which data must be collected and pooled over months (Azadi et al., n.d.; Rajalingham et al., 2021). Moreover, compared with acute methods such as using optical fibers or direct illumination, a chronically implantable light delivery device reduces the risk of tissue damage as well as the risk of infection from open cranial chambers. In this article we explain the surgical techniques and procedures for Opto-Array implantation in macaque monkeys. We typically perform two separate surgeries; in the first surgery, we perform virus injection using an injector array. After a period of 4 to 8 weeks, the second surgery takes place, in which we confirm virus expression and implant the Opto-Array.

## 2. Methods

### 2.1. First surgery: virus injection

#### 2.1.1. Head stabilization

We use a low-profile head holder (Jerry-Rig USA, item #1) to stabilize the animal’s head (Figure 1). For secure head placement, the ridges on the tooth bar should be placed in between the molar teeth symmetrically, and the teeth should lay flat on the tooth bar. The orbit bar should be secured in a horizontal position, parallel to the base. Unlike stereotaxic devices, this head holder does not require earbars, making it possible to fully retract temporalis muscle. Moreover, its rotational ball technology provides the ability to rotate the head during the surgery.

**Figure 1:**
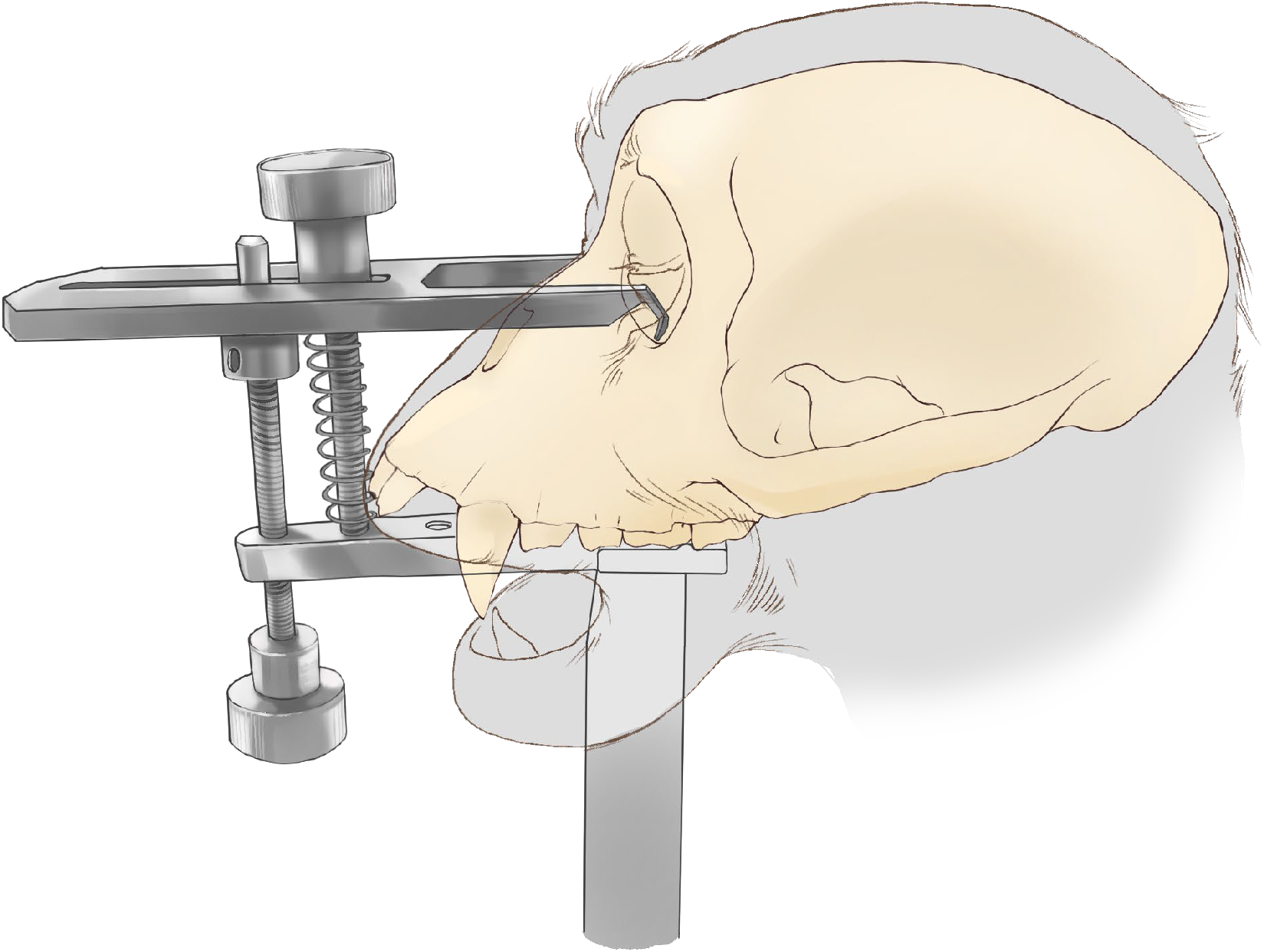
head placement in low-profile head holder.

#### 2.1.2. Incision

We perform a long coronal incision on the skin crossing zygomatic arches on both sides (Figure 2) using a surgical blade (#15). This provides access to the temporal and zygomatic bones, as well as parietal bones on which we will implant the pedestal connector. After separating the skin margin from the fascia, we perform the fascia incision parallel to the skin margin with the same blade. It is important to only cut the layers of the fascia and not the epimysium of the underlying temporalis muscle. Alternatively, the incision through the fascia can be made using Mayo scissors to avoid any potential damage to the muscle capsule.

**Figure 2:**
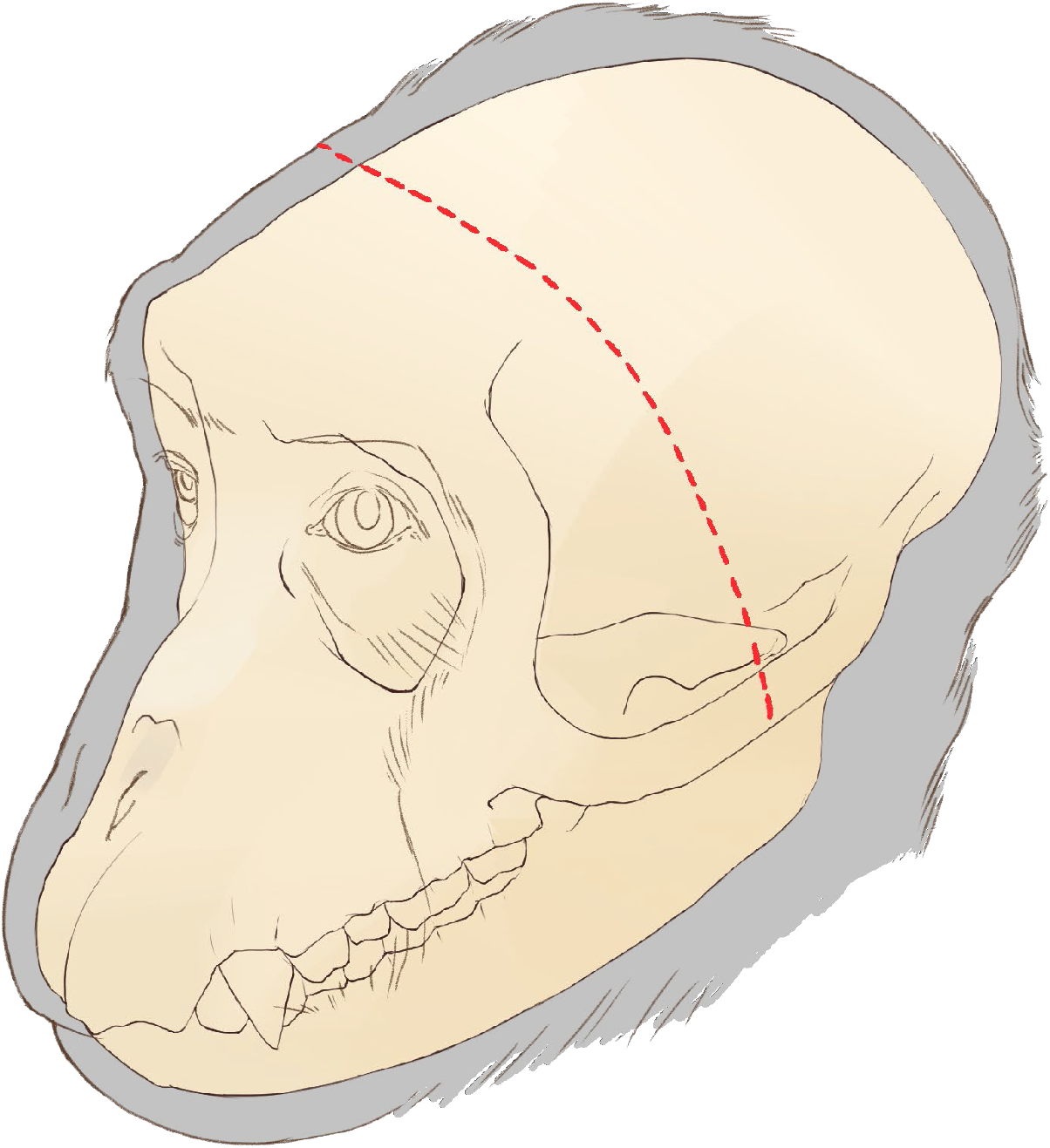
coronal incision

#### 2.1.3. Skull exposure

Removing the zygomatic arch allows us to fully reflect temporalis muscle and expose the temporal bone. To expose the arch, make a 10 - 15 mm horizontal incision in the temporalis muscle fascia covering the center of the arch. Use a periosteal elevator to detach the muscle and fascia attachments from the arch. These attachments are mainly temporalis muscle and partly masseter and zygomaticus major muscles. After isolating the arch, cut both ends using rongeurs and extract the arch (Figure 3). Use rongeurs to remove all the remaining parts of the arch on the zygomatic bone and temporal bone and smooth any sharp edges. Bone wax can be used to stop any osteal bleeding. Close the fascia incision initially made over the arch with an absorbable suture (e.g., vicryl 3-0).

**Figure 3:**
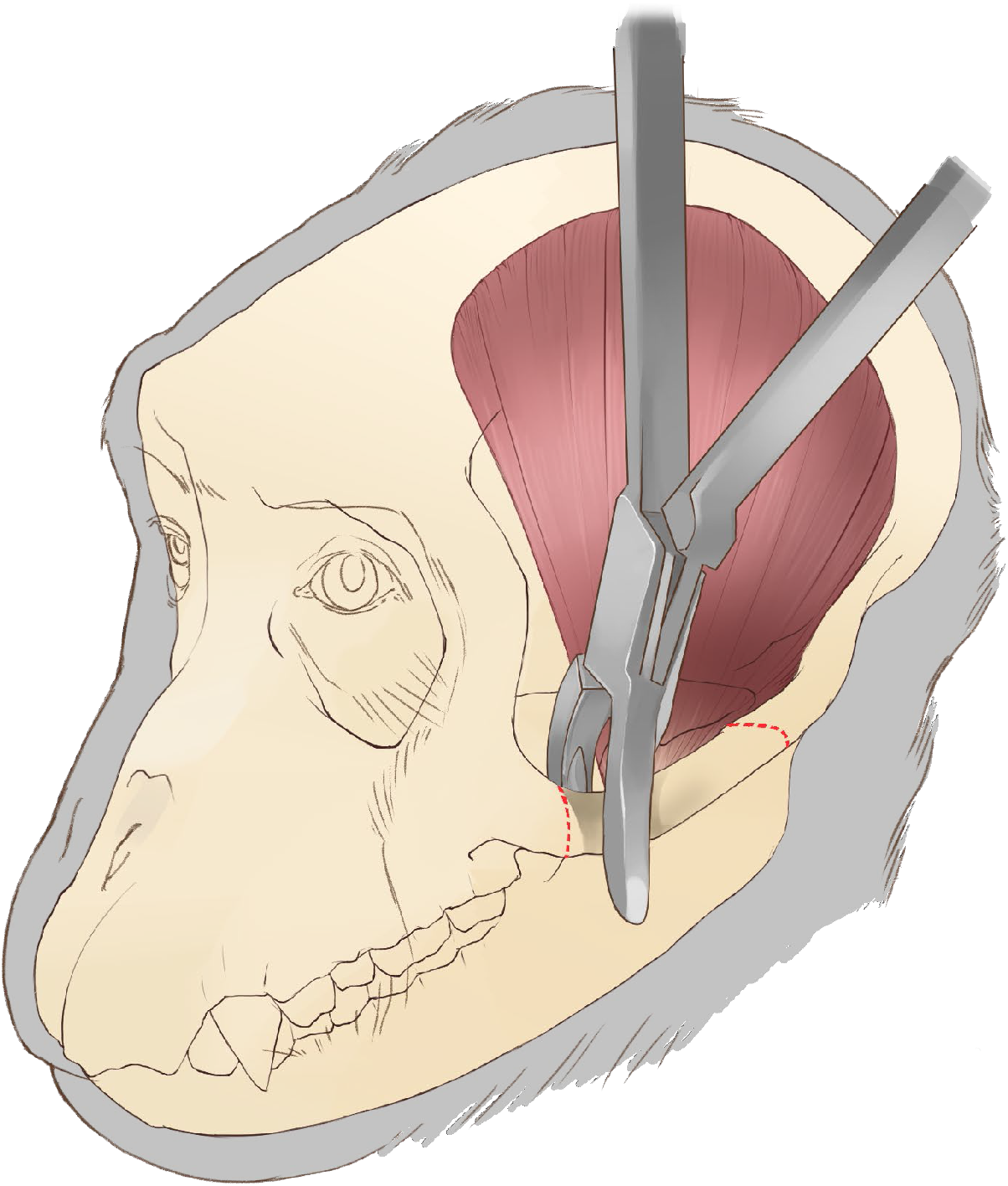
removing the zygomatic arch

The temporal bone can be exposed by detaching the temporalis muscle from its origins on the dorsal aspect of the skull using a periosteal elevator to reflect it towards its ventral insertions. Use traction on the temporalis muscle to maximize access to the temporal bone (Figure 4).

**Figure 4:**
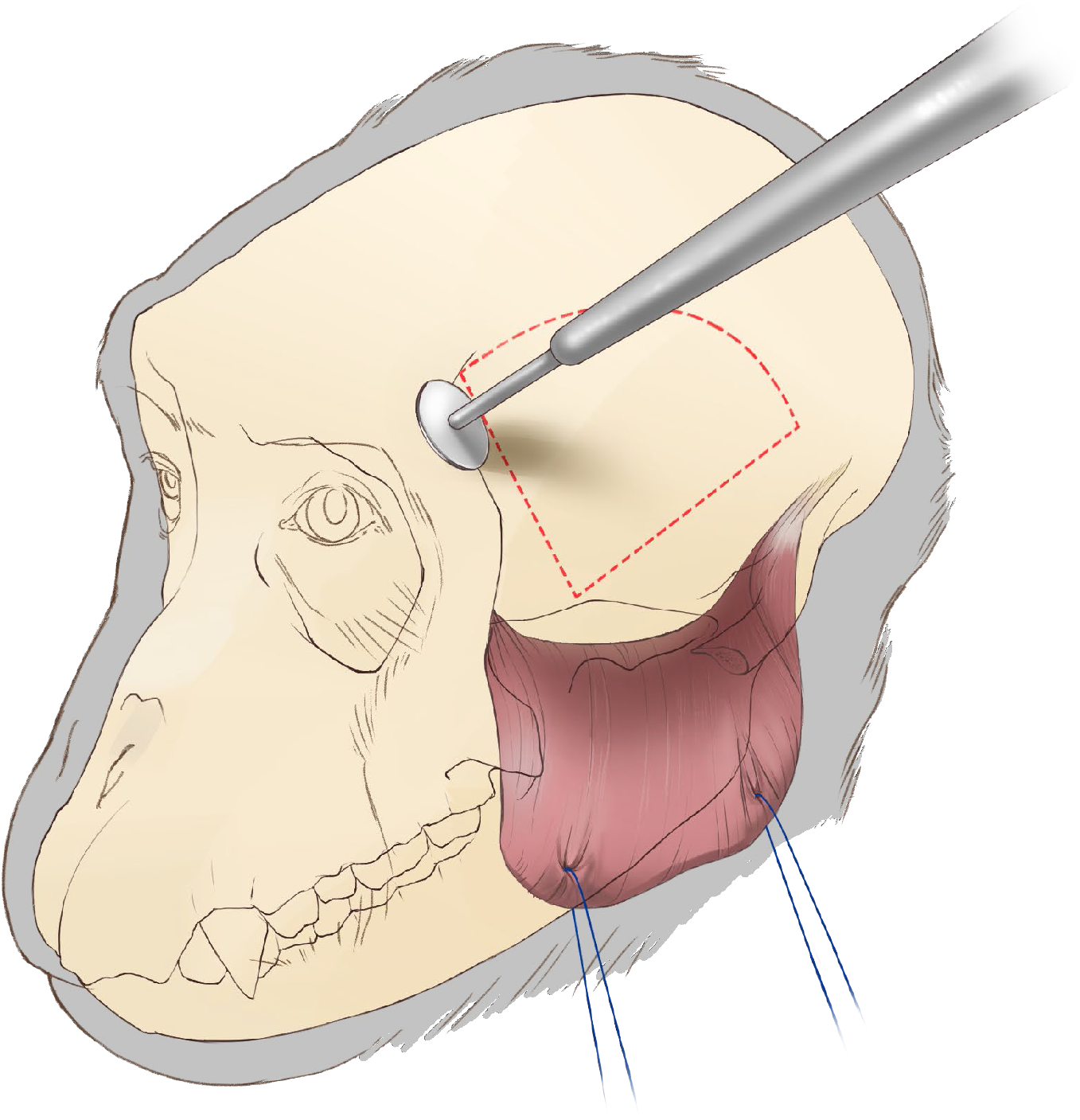
craniotomy with circular cutting wheel

#### 2.1.4. Craniotomy (bone flap removal)

For the bone flap, use a circular cutting wheel (Figure 4; item #2) at a 45° angle to the surface of the bone. This is important since later we replace the bone flap for closure. To prevent dura damage during craniotomy, try not to drill through the full thickness of the skull, instead finish the opening by using two periosteal elevators to break the bone flap free from the cranium. Alternatively (less preferred method), the craniotomy can be performed using drill bits (item #3) and covered by titanium straps (item #4) or teflon (item #5) for closure. While removing the bone flap, carefully detach any dura adhesions by sliding a second elevator between the flap and the dura.

After removing the bone flap, carefully detach any dura adhesions to the skull around the craniotomy margin. Special caution is needed near the ventral posterior aspect, near the interaural canal, where the vein of Labbé runs from the dura to the bone. Smooth the dorsal and anterior margins of the craniotomy and the bone flap edges using rongeurs. Using a drill bit (item #2), drill four holes on the dorsal edge of the craniotomy which later will be used for suturing the bone flap back (Figure 5). During drilling, use a ‘surgical spoon’ (item #6) between the bone and dura to protect the dura and the cortex. Drill these holes at angle - exit hole closer to the bone margin than the entry hole - to make it easier to pass a needle through.

**Figure 5:**
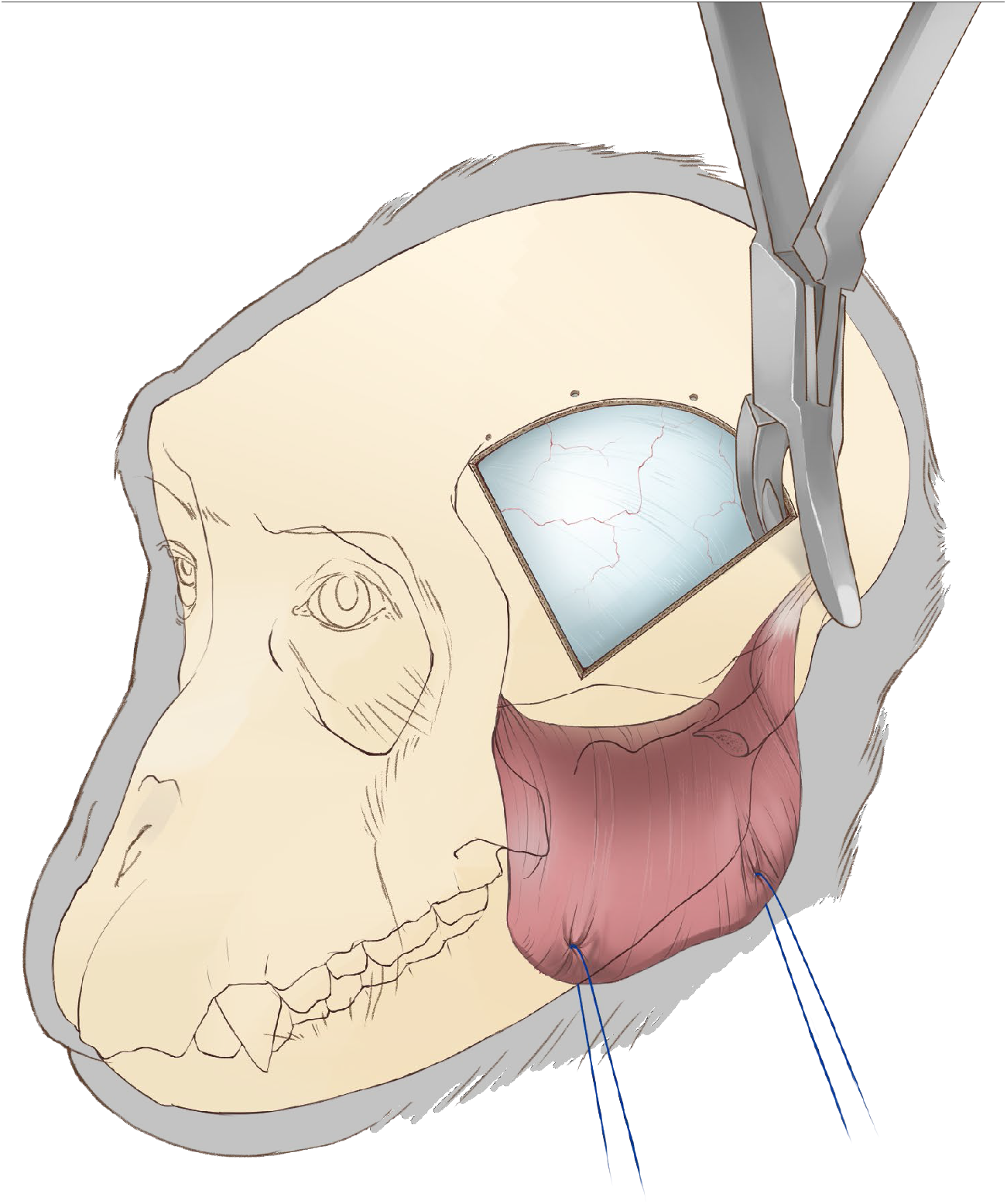
expanding the craniotomy

Expand the ventral aspect of the craniotomy by rongeuring the temporal and sphenoid bones to the base skull (Figure 5). Use an elevator to ensure the bone is separated from the dura before rongeuring to avoid tearing the dura or bridging vessels. This expansion of the craniotomy is crucial for accessing the lateral aspect of inferior temporal cortex. Bone wax can be used to control osteal bleeding.

#### 2.1.5. Dural opening

A T-shape incision in the dura provides the best access to central and anterior IT (Figure 6). Start the dura opening with a dorsal curved incision with approximately a 4 mm margin from the edge of the craniotomy, followed by a right-angled incision towards the temporal pole reaching the rostral-ventral corner of the craniotomy. Two suture lines attached to small hemostatic forceps can be used to provide traction.

**Figure 6:**
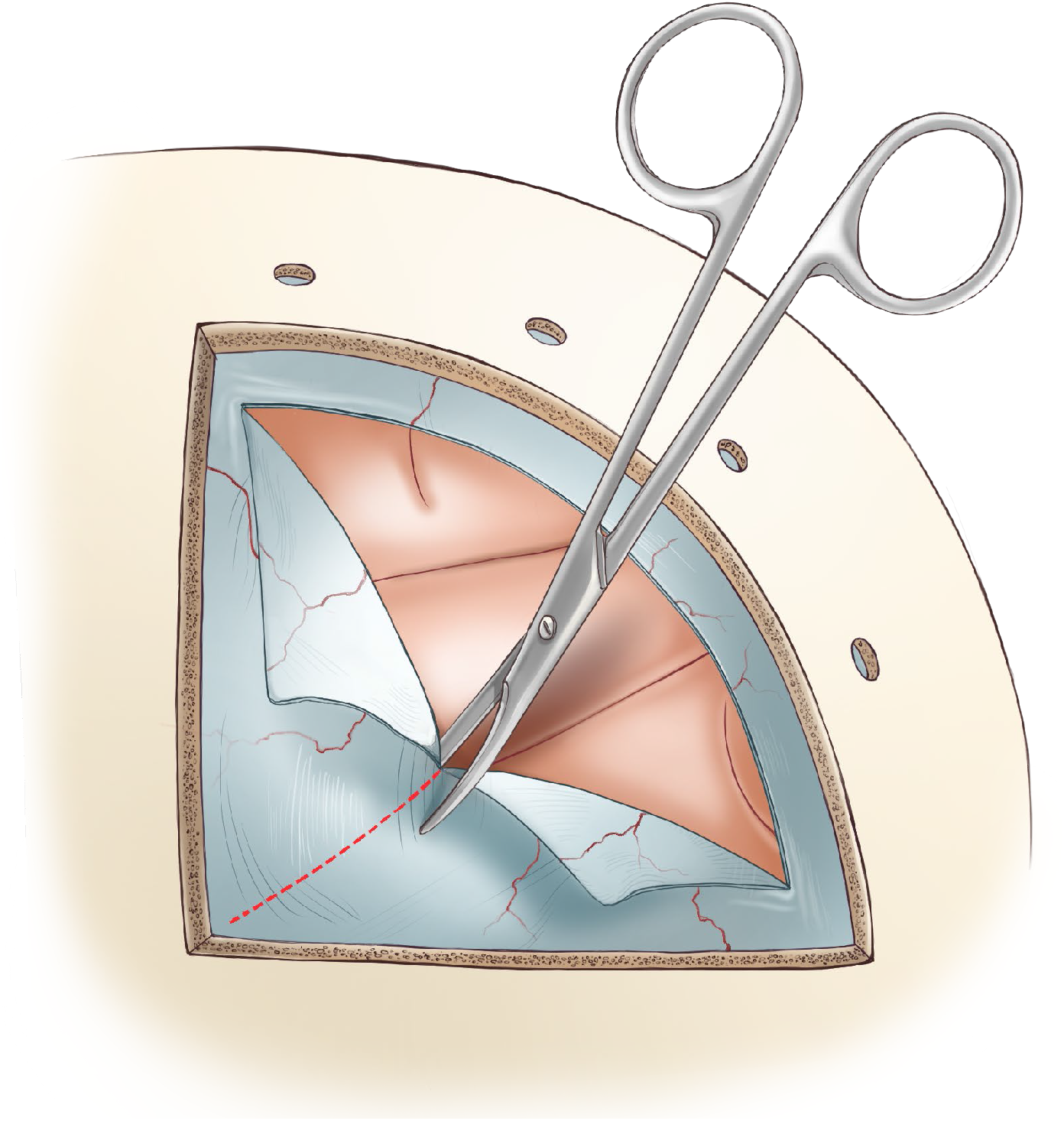
dural opening

#### 2.1.6. Virus injection

To speed up the virus injection procedure, use an injector array allowing efficient and uniform vector delivery with four channels in parallel (Fredericks et al., 2020) (Figure 7). Covering the lateral surface of central IT (the area of interest of this paper) often requires 3 to 4 placements of the injection array, injecting 10 μl per needle at a 0.5 μl/min rate, resulting in 40 μl per array placement in 20 minutes. At the end of each injection, leave a 10-minute wait time to allow the injected liquid to disperse and diffuse into the cortical tissue before removing the needles.

**Figure 7:**
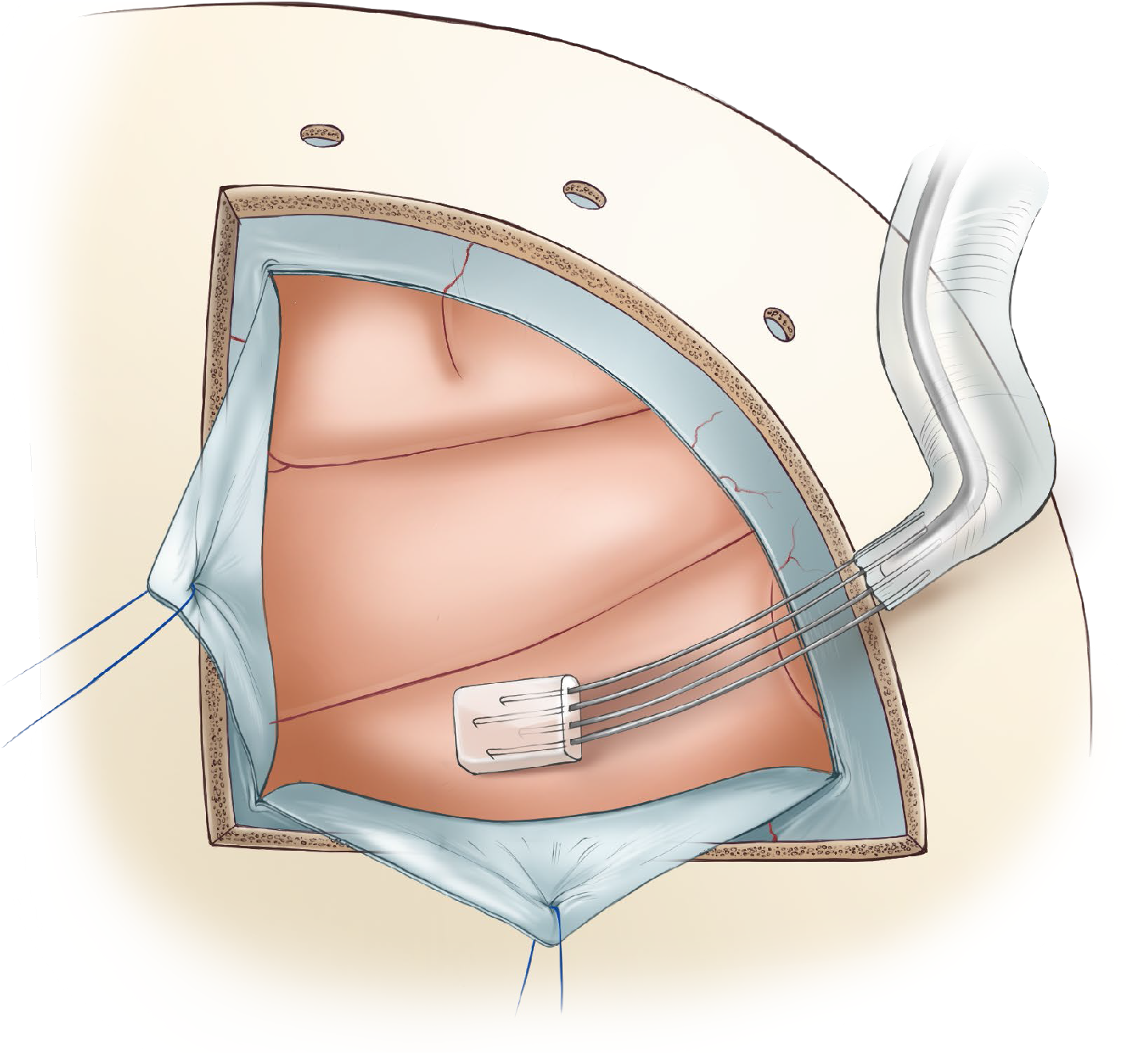
virus injection

After the injection procedure, place sterile transparent film (item #7) on the pial surface and mark the injection site, the craniotomy margins, and other anatomical landmarks, then remove the film. This film will be used to pinpoint the injection site in the second surgery to make a proper dural opening.

#### 2.1.7. Dural closure

Before closing the dura place a piece of artificial dura (item #8) between the dura and the cortex to prevent any potential adhesions prior to the second surgery. Another piece of artificial dura can also be placed between the dura and the bone flap before closing the craniotomy.

Use absorbable suture lines (vicryl 5-0) for dural closure. First, secure the tips of each dural flap to one another, and then attach the joined region to the dorsal edge of the dura opening with single interrupted sutures. Close the dorsal-ventral incision with a simple continuous suture pattern with 1-2 mm spacing, starting from the rostral-ventral corner of the craniotomy. The dura closure procedure will be completed by closing both sides of the dorsal incision (Figure 8).

**Figure 8:**
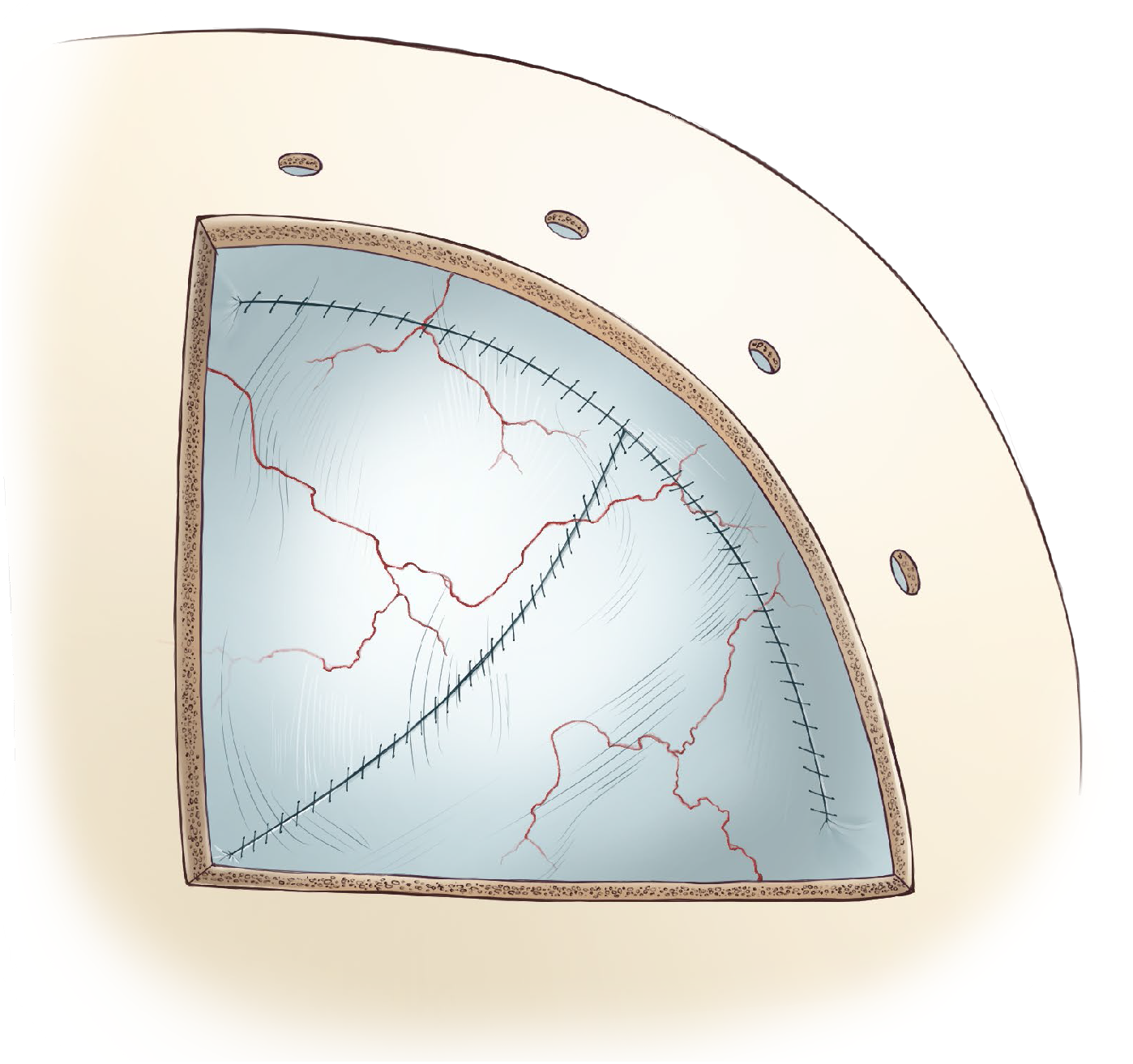
dural closure after virus injection

#### 2.1.8. Bone flap replacement

Use absorbable sutures for this procedure (vicryl 3-0). First, pass the suture lines through each of the holes previously drilled into the skull. Place a grooved surgical spoon (item #6) under the skull while passing the needles through the holes to protect the dura and cortex. Pass the suture lines through corresponding holes in the bone flap. Finally, fix the bone flap with square knots (two half knots in opposite directions) allowing gradual tightening followed by two more half knots to secure. Optionally, injectable bone putty can be used to fill the gaps (item #9) and promote osteoinductivity.

#### 2.1.9. Closing

Replace the origin of the temporalis muscle and suture back in place; this often requires two or three sutures on the frontal and occipital sides. Close the fascia with a simple interrupted or continuous suture pattern, and the skin with simple interrupted or continuous horizontal mattress suture patterns. Use absorbable suture line (vicryl 3-0) for the muscle and fascia and non-absorbable (polyester 3-0) for the skin.

### 2.2. Second surgery - Opto-Array implantation

The second surgery is performed 4 to 8 weeks after the first surgery. The goal of the second surgery is to confirm virus expression and implant the Opto-Array. The general approach and closing is the same as the first surgery. Drill the bone flap on the same lines as the first surgery if the bone flap has fused with the skull.

#### 2.1.1. Dural opening

After removing the external artificial dura (if used) between the dura and skull,use the transparent film marked in the first surgery to find the injected site. Perform a horizontal dural incision parallel to the base skull at least 10 mm dorsal to the injection site (Figure 9), followed by two parallel incisions extending to the ventral margin of the craniotomy. The distance between the parallel incisions should be4 mm, which is the distance between the Opto-Array holes. After opening the dura, remove the artificial dura placed between the dura and cortex in the first surgery, to expose the cortex.

**Figure 9:**
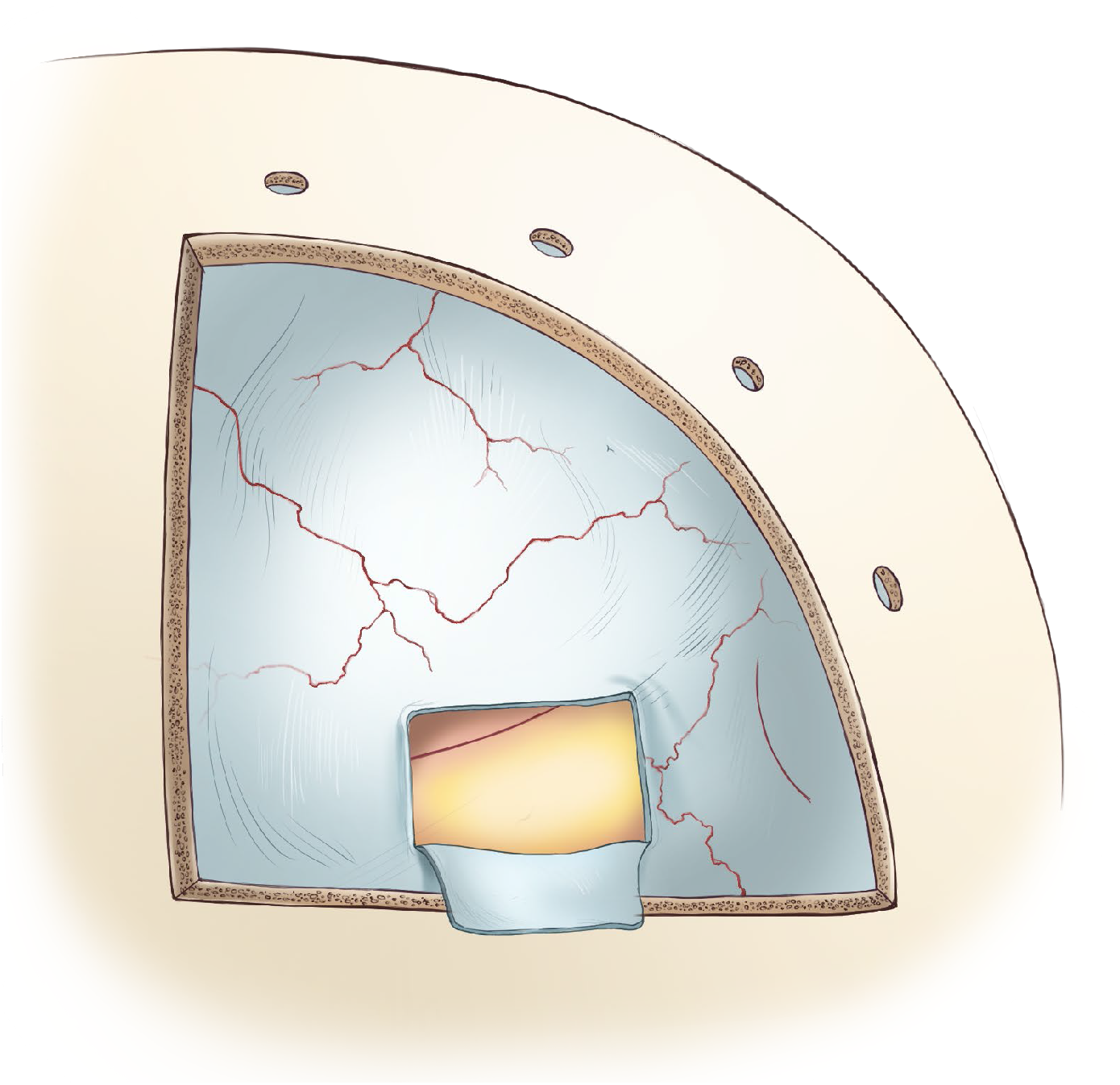
verification of virus expression

#### 2.2.2. Virus expression verification

To confirm expression, check and document the fluorescent signature of the virus. Use a light source and filter with wavelengths matching the virus fluorescent protein (for example, 440–460-nm excitation light, 500-nm longpass filter for GFP).

#### 2.2.3. Opto-Array implantation

The array connector and cables need to be gas sterilized prior to the surgery. Before implanting the array, first, secure the connector pedestal to the skull with two self-drilling titanium screws (6 mm long; item #10). The remaining screws to secure the pedestal connector will be placed later in the procedure.

Implant the array by suturing the holes located on its four corners to the edges of the dural window that was opened in the previous step (Figure 10). Surgeon’s knots of braided non-absorbable suture should be used to secure the array in place (5-0 silk).

**Figure 10:**
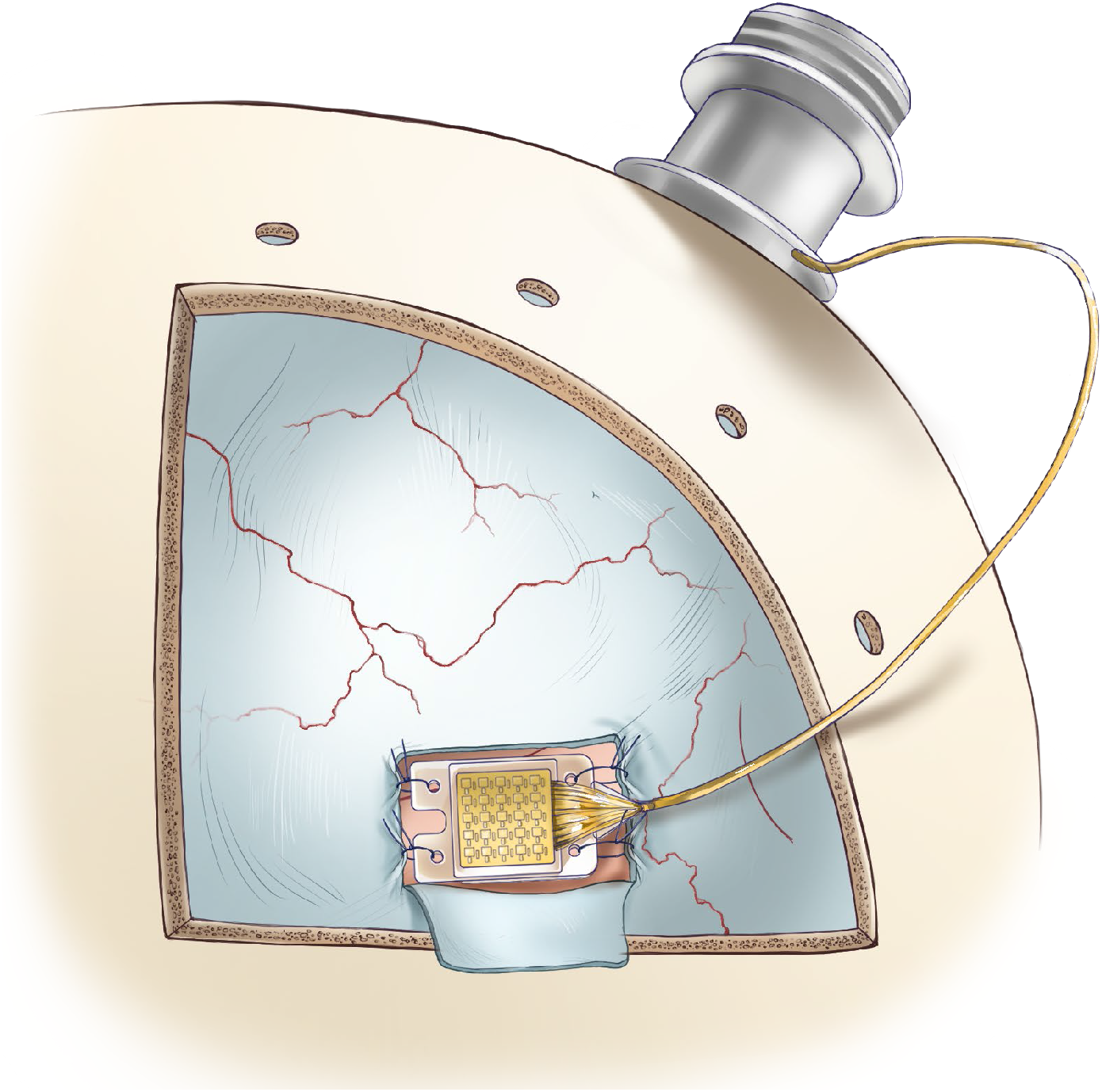
Opto-Array implantation

The skin around the pedestal implant may gradually recede, exposing the wires. To protect the wires, cover them with a medical-grade polypropylene mesh (item #11), made for hernioplasty surgeries. Secure the wire and the mesh on the skull with titanium plates (item #12) and self-drilling titanium screws (3.5 mm long; item #13).

Test the Opto-Array to make sure it has not been damaged during the array and wire implantation. At this stage, the array can still be replaced if necessary, since the pedestal connector has not been permanently implanted.

#### 2.2.4. Dural closure

Place the dura flap back to cover the Opto-Array, suture the dorsal edge followed by the anterior edge. For dura closure, use braided absorbable sutures (vicryl 5-0) in a simple interrupted pattern. It is important not to over tighten the dura flap on the array in order to avoid cortical damage. After closure, cover the dura with an artificial collagen-based dural graft (item #14) to conceal any gap remaining in the dural opening and also to facilitate dural cell growth in the region of the opening.

#### 2.2.5. Pedestal connector implantation

After closing the dura and replacing the bone flap (per 2.1.8), secure the pedestal with self-drilling titanium screws (6 mm long; item #10). However, self-drilling screws may not secure the pedestal in the long term. Thus, one option is to use self-tapping titanium screws (item #15) instead of the self-drilling screws, or the implant can be reinforced using a dental adhesive cement system applied on the skull and under the pedestal connector. Moreover, two or three self tapping titanium screws (item #16) can be implanted around the pedestal, and covered with dental acrylic contiguous with the acrylic covering the base of pedestal. The dental adhesive cement system (item #17) can also be applied to fill the gap between the base of the pedestal and the skull before using acrylic.

All procedures presented in this manuscript were conducted under an Animal Study Protocol approved by the National Institute of Mental Health Animal Care and Use Committee.

### 2.3. Behavioral Experiment Methods

As a proof of concept, we measured the behavioral performance of a rhesus monkey (*macaca mulatta*) trained to detect cortical stimulation using an Opto-Array implanted on IT cortex of the left hemisphere. In the first surgery we injected Adeno Associated Virus 5 (AAV5) expressing excitatory opsin C1V1 under the CaMKIIa promoter. In the second surgery we confirmed virus expression via fluorescent signature and implanted an Opto-Array on central IT cortex. Following recovery from the second surgery, we trained the animal to behaviorally detect the optogenetic cortical stimulation evoked by illumination of a single LED on the Opto-Array (Figure 13.A). We used a psychophysical detection task in which the animal was rewarded for correctly identifying whether a given trial contained optogenetic stimulation (Azadi et al., n.d.; Rajalingham et al., 2021). Each trial started with the animal fixating on a central fixation point (black-on-white bullseye, 0.4° outer diameter and 0.2° inner diameter) for 500 ms. Then an image appeared on the screen; for1000 ms while the animal was required to fixate on the central fixation point. In 50% of the trials (randomly selected), 500 ms after the image appeared, a cortical stimulation pulse was delivered by illuminating an LED. The stimulation intensities were randomly chosen from three stimulation powers: 0.4, 1.2, 2.5 mW, and three stimulation durations:100ms, 200ms, and 400ms. The choice phase was cued by the disappearance of the fixation point and the image, after which two visual targets were presented on the vertical midline of the screen (at 5 deg eccentricity). The animal received a liquid reward for saccading to a response target associated with the trial condition (“stimulation” or “no-stimulation”).

**Figure 11:**
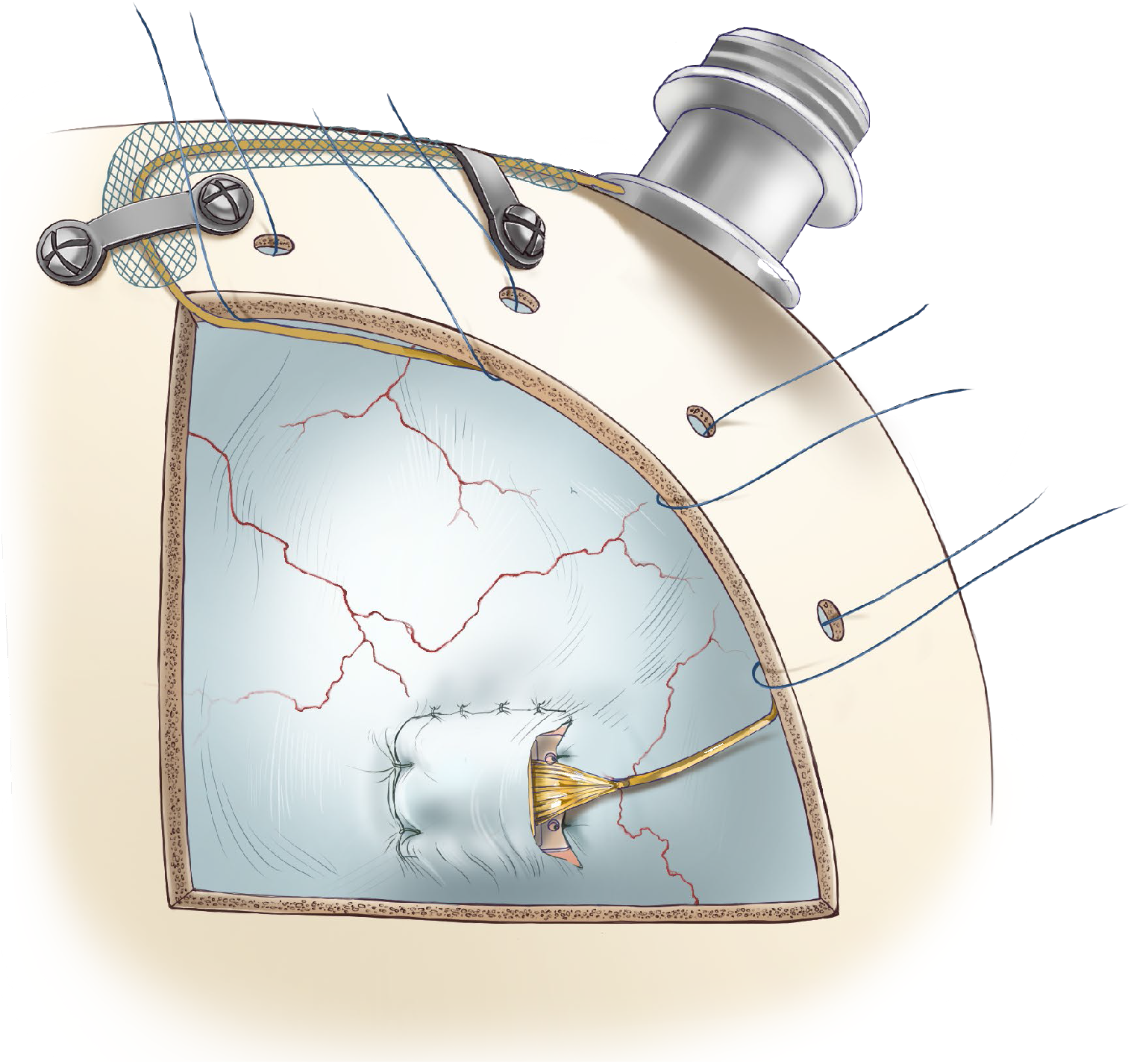
dural closure after Opto-Array implantation

**Figure 12:**
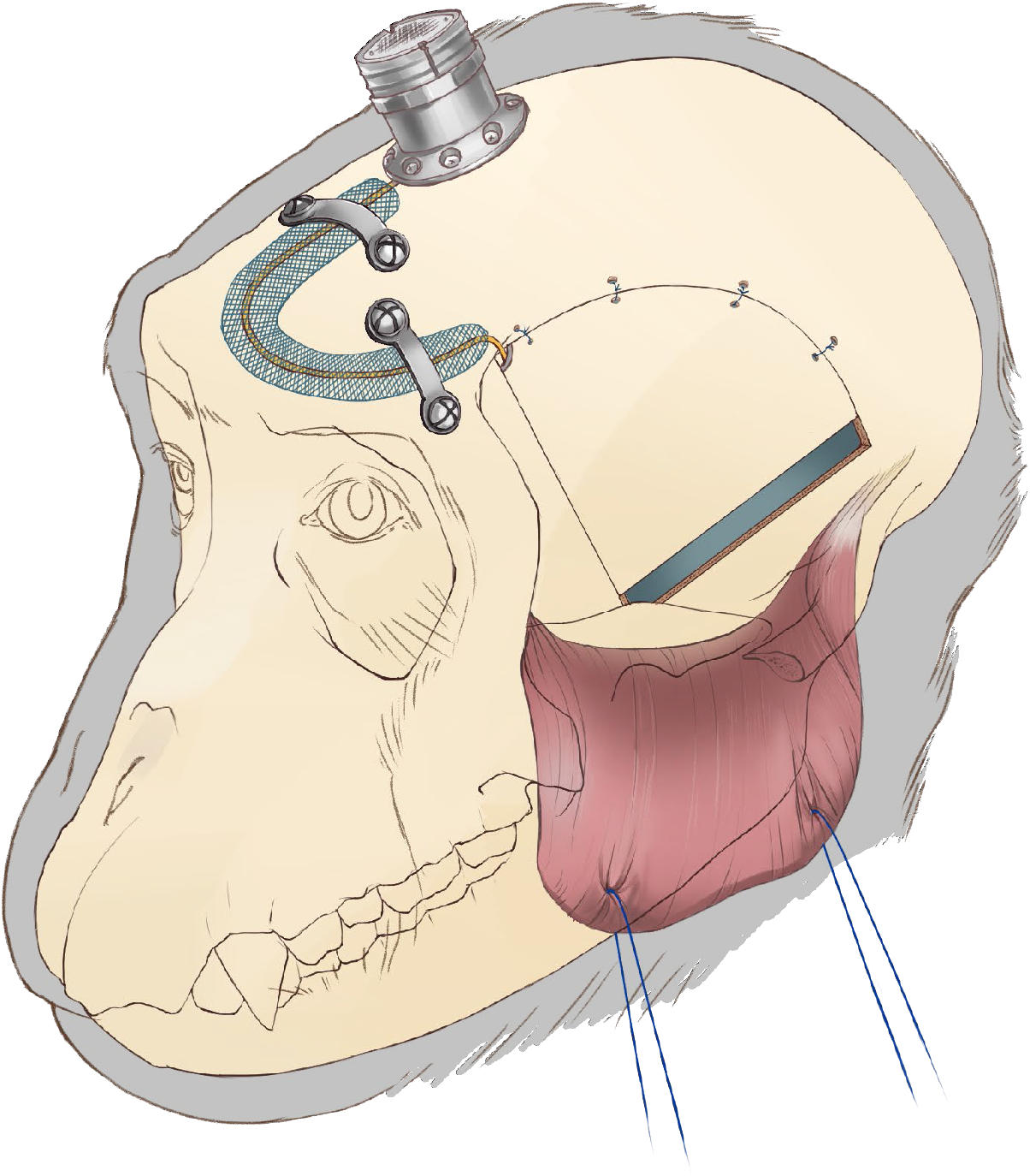
implantation of the pedestal connector and securing the wire

**Figure 13:**
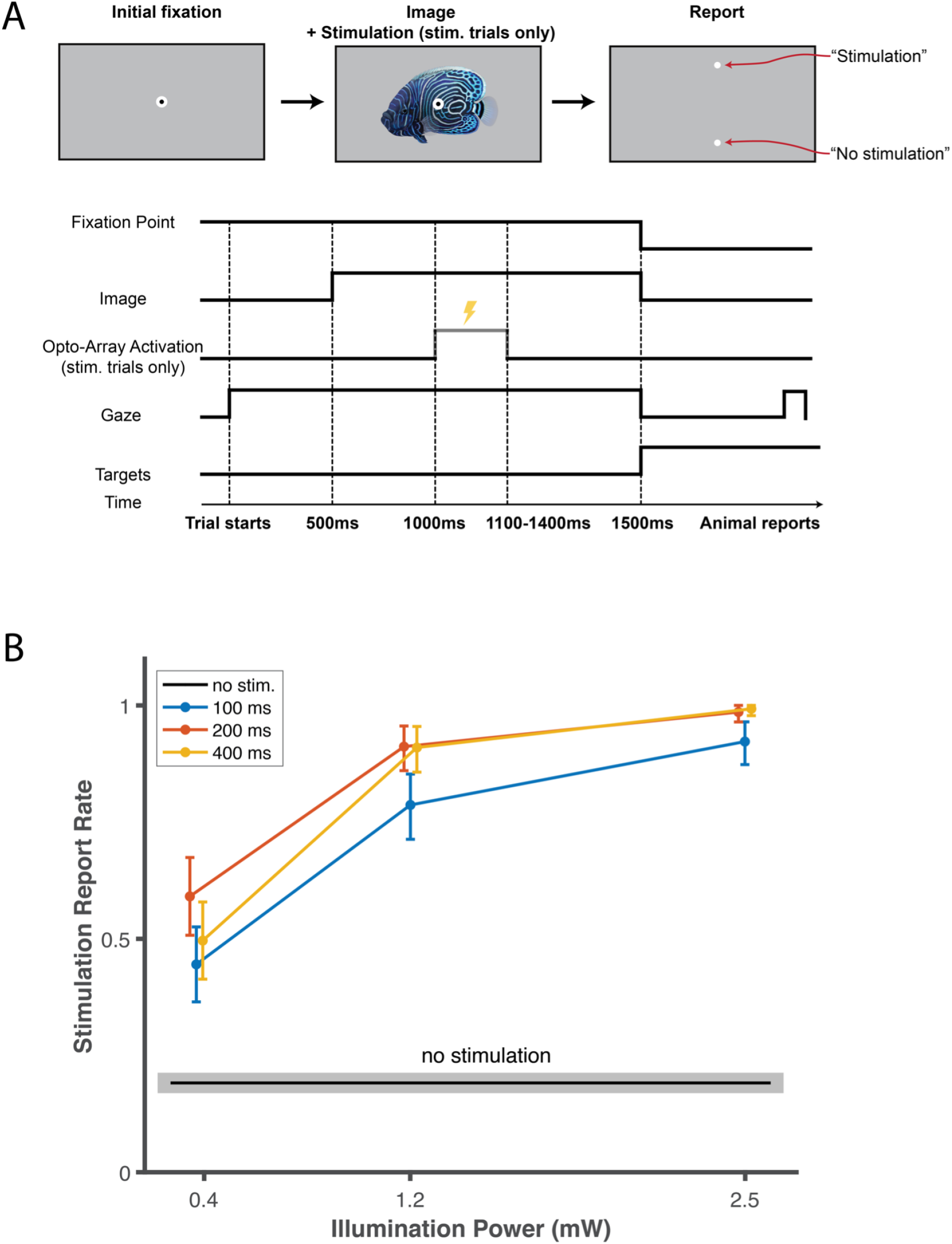
Behavioral detectability of Opto-Array cortical stimulation at different illumination duration and powers. A. Schematic illustration of the behavioral task. The monkey’s task was to report whether a given trial did or did not include cortical stimulation. After 500 ms of initial central fixation an image appeared on the screen. The animal was required to keep fixation for 1000 ms until the fixation point and the image disappeared. In “stimulation” trials (randomly selected 50% of trials) we stimulated the cortex by activating the Opto-Array. The activations varied in duration (100, 200, and 400 ms) and light power 0.4, 1.2, and 2.5 mW). The stimulation (when present) began 1000ms after the animal started fixation (500ms after image appearance) After the disappearance of the image from the screen, two response targets appeared (each corresponding to a trial-type; “stimulation” and “no stimulation”), and the monkey reported a choice by fixating on one response target. A liquid reward was delivered if the choice was correct. B. Behavioral performance as a function of duration -and power of optogenetic stimulation in the detection task. The y-axis is the monkey’s stimulation report rate, and the x-axis is illumination power. The blue, red and yellow colors, respectively, represent stimulation trials with 100, 200 and 400 ms of illumination duration (hit rates). The black line represents data from no stimulation trials (false alarm rate). The error bars and the shaded gray area represent bootstrapped 95% confidence intervals (with 10,000 resamples).

## 3. Results

The animal was able to detect the optogenetic impulse across all tested illumination powers and durations (*n* = 2416 trials; permutation test, *p* < 0.001 for all comparisons, benjamini-hochberg corrected; Figure 13.B). For each stimulation duration, the stimulation was detected at a higher rate when the light intensity was higher (Spearman’s correlation between performance (hit rate) and illumination power *r* = 0.52, 0.60, 0.58, *p* < 0.001 respectively for 100ms, 200ms, 400ms illumination durations). To summarize the results and analyze the effect of illumination duration and power, we ran the following linear regression on the data:

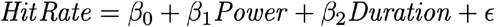

Where *Power* and *Duration* of illumination are independent variables and *HitRate* is the animal performance for detecting optogenetic stimulation. The results showed this regression model can significantly explain the performance of the animal (*R*^*2*^ = 0.42, *F*(3, 2412) = 848, *p* < 0.001). Moreover, both power and duration of illumination significantly modulate the performance (*β*_*0*_ = 0.20, *t*(2412) = 18.9, *p* < 0.001; *β*_*1*_ = 0.22, *t*(2412) = 19.3, *p* < 0.001; *β*_*2*_ = 0.02, *t*(2412) = 13.2, p < 0.001).

## 4. Discussion

In this study, we first documented a procedure for virus injection and implantation of the Opto-Array on IT cortex of macaque monkeys. We then presented results from a stimulation detection task to validate the functionality of the Opto-Array under different power and durations of illumination. These results demonstrate that the surgical procedure described in this paper creates the conditions for eliciting reliable behavioral effects from a range of optogenetic illumination intensities and durations. The similarity of performance across different stimulation durations suggests that illumination longer than 200 ms may be reaching a “ceiling effect” with respect to the channel kinetics of adaptation for the optogenetic channel we used, C1V1. Notably, to achieve these behavioral effects, the illumination powers used in the present experiment were far below the maximum capacity of the Opto-Array. In this experiment, we used a single LED powered to beween 1% and 4% of the maximum individual light capacity of a single LED. Other experimental conditions (for example, a less sensitive behavioral task, or usage of an inhibitory opsin) may require the activation of more LEDs on the array and/or a higher power of activation.

The Opto-Array allows us to harness the advantages of optogenetic methods such as cell targeting, and stimulation and inhibition of neuronal activity in NHPs. Using this chronically implantable array of LEDs also facilitates uniform, replicable stimulation and/or inhibition across multiple sessions. Moreover, the chronic nature of this array does not require a cranial chamber thereby reducing the risk of infection and tissue damage. In Table 1, we list the benefits and limitations of the Opto-Array over other methods. Here we also presented a detailed step-by-step surgical protocol for virus injections and implantation of the Opto-Array on IT cortex of macaque monkeys. In Table 2, we list some important and novel techniques and surgical approaches presented in this manuscript. Moreover, the surgical approach presented here can be applied to other procedures such as implanting chronic electrode arrays.

**Table 1:** comparison of different tools and methods used for perturbation of neuronal activity.

|  | Acute microelectrodes | Chronically implantable microelectrodes | Optic fiber | Opto-Array |
| --- | --- | --- | --- | --- |
| Stimulation | yes | yes | yes | yes |
| Inhibition | no | no | yes | yes |
| Cell type targeting | no | no | yes | yes |
| Perturbation of the same site over multiple sessions | no | yes | no | yes |
| Suitable for cortical surface | yes | yes | yes | yes |
| Suitable for deep brain structures | yes | yes | yes | no |
| Necessity of open cranial chambers | yes | no | yes | no |
| Damage caused by penetration | yes | yes | yes | no |

**Table 2:** list of some important and novel techniques and surgical approaches presented in this manuscript.

| Commonly used | Presented in this paper | Advantages |
| --- | --- | --- |
| Stereotaxic systems | A low profile head-holder | Better access to the ventral areas since it does not require usage of earbars; allowing rotation of the head during the surgery. |
| Cutting through temporalis muscle | Reflecting temporalis muscle | Less bleeding and tissue damage; dramatically shortens the duration of the surgery; increasing the field of exposure; all above resulting in a faster healing process. |
| Single needle for virus injection | An injector array | Efficient and uniform vector delivery in parallel; resulting in a shorter procedure. |
| Artificial dura or dural grafts for dural closure | Suturing back reflected dura | Assist the healing process of the dura; isolate the brain; decreasing the risk of infection. |
| Burrs for craniotomy | Circular cutting wheel | Minimizing the gap between the bone flap and craniotomy, allowing bone flap replacement. |
| Titanium or teflon implants to cover the craniotomy | Bone flap replacement | Minimizing the usage of foreign bodies; facilitating the healing process; decreasing the chance of complications and infections. |
| Self-drilling screws for implanting the pedestal connector | Dental adhesive cement system under the pedestal to fill the gap between the base of pedestal and skull | Decreasing the chance of local infection and failure by minimizing the empty space under the implant. |
| Leaving the wire under the skin without any protection | Medical-grade polypropylene mesh to secure the wire | Protecting the wire in case of skin recession. |

Surgical access to ventral and ventrolateral surface cortex, e.g. IT cortex, in nonhuman primates can be technically challenging, and the lack of explicit and detailed protocols in the literature makes replication of these procedures difficult. We have provided a detailed description of one method for targeting virus and LED arrays to ventrolateral surface cortex; in doing so, we hope to improve the replicability and reliability of optogenetic experiments in nonhuman primates.

## Acknowledgements

We thank Erina He from NIH Medical Arts and Emily Lopez for their critical help in preparing the illustrations; This research was supported by the Intramural Research Program of the NIMH ZIAMH002958 (to A.A.).

## Author contributions

R.A. prepared the illustrations and the original manuscript with guidance from M.E. and A.A.; S.B. ran the experiment and collected the behavioral data. S.B. and R.A. analyzed the data and prepared the results. S.B., M.E. and A.A. edited the manuscript; all authors reviewed the manuscript.

## Competing interests

The authors declare that there is no conflict of interest.

---

**Table S1:** list of items

| Item # | Item name | Commercial name | Company | Catalog number | Webpage |
| --- | --- | --- | --- | --- | --- |
| 1 | Head holder | LOW PROFILE:<br>Stainless Steel<br>Holding Instrument | Jerry-Rig USA | JR-SSHC | http://jerry-rig.com/medical/ |
| 2 | Circular saw | Cutting wheels | Pfingst &<br>Company | H041047 (S)<br>H041061 (M)<br>H041073 (L) | https://www.pfingstco.com/products/horico-diamonds-rotary-instruments/041/ |
| 3 | Drill bit | Drill Bits | Roboz Surgical<br>Instrument<br>Company | RS-6280C-2 (#2)<br>RS-6280C-5 (#5)<br>RS-6280C-8 (#8) | http://shopping.roboz.com/Surgical-Instrument-Online-Shopping?search=RS6280 |
| 4 | Titanium<br>straps | Low Profile Neuro<br>Plate | KLS Martin LP | 25-316-01-09 | https://www.thomasrecording.com/titanium-mini-plate-system |
| 5 | Teflon | HDPE Sheet<br>1/16" x 12" x 24" | United States<br>Plastic<br>Corporation | 46603 | https://www.usplastic.com/catalog/item.aspx?itemid=132518&gclid=CjwKCAiAz--OBhBIEiwAG1rIOlkp4ro-mlVTERnkWq-WWdeqtkytM9JEzJGHppDLWtldhIKIyyHbxoCjQ4QAvD_BwE |
| 6 | Surgical<br>spoons | Stainless Steel<br>Surgical Spoons | Jerry-Rig USA | JR-SSSS | http://jerry-rig.com/medical/ |
| 7 | Transparent<br>film | Uncoated<br>Transparency Film | Apollo | 829911 | https://www.staples.com/Apollo-Plain-Paper-Copier-Transparency-Film-Clear-8-1-2-W-x-11-H-100-Box/product_829911 |
| 8 | Artificial<br>dura | Tecoflex EG-93A<br>Thickness 0.005" | Lubrizol | NA | https://www.lubrizol.com/Health/Medical/Polymers/Tecoflex-TPU |
| 9 | Injectable | ALLOMATRIX | WRIGHT | 86000100 | https://www.wright.com/products-biologics/allomatrix- |
|  | bone pottey |  |  |  | injectable-dbm-paste |
| 10 | Self drilling 6mm screw | Level one cmf screw, maxdrive, drill free size:1.5 X 6 mm, MATL:TI-6AL-4V | KLS Martin LP | 25-878-06-09<br>10888118047707 | https://www.klsmartin.com/fileadmin/user_upload/Homepage/Mediathek/90-440-02-04_LevelOne_1.0_Micro.pdf |
| 11 | Hernia repair mesh | Medtronic Duatene, bilayer mesh monofilament polypropylene | Medtronic | DTN1610E | https://www.medtronic.com/animal-health/en-us/products/hernia-repair/duatene-bilayer-mesh.html |
| 12 | Titanium plate | Level one neuro plate, low profile, str, neuro screw size:2 hole, 15 mm, T=0.6 mm, MATL:CP TITANIUM | KLS Martin LP | 25-302-13-09<br>00888118037732 | https://www.klsmartin.com/fileadmin/user_upload/Homepage/Mediathek/90-343-02_Low_Profile_Neuro_System.pdf |
| 13 | Self drilling 3.5mm screw | Level one neuro screw, maxdrive, drill free size:1.5 X 3.5 mm, MATL:TI-6AL-4V | KLS Martin LP | 25-975-03-09<br>10888118048742 | https://www.klsmartin.com/fileadmin/user_upload/Homepage/Mediathek/90-440-02-04_LevelOne_1.0_Micro.pdf |
| 14 | Dural graft | DuraGen Plus | INTEGRA | NA | https://www.integralife.com/duragen-plus/product/dural-repair-grafts-duragen-plus |
| 15 | Titanium screw- self tapping | 2.0mm Cortical Screw, Self-tap, Hex, Titanium | Movora | T200.06T | https://usstore.movora.com/products/2-0mm-cortical-screw-self-tap-hex-titanium.html |
| 16 | Titanium screw- self | 2.7mm Cortical Screw, Self-tap, | Movora | T270.06T | https://usstore.movora.com/products/2-7mm-cortical-screw-with-2-5mm-head-recess-non-self-tap-hex- |
|  | tapping | Hex, Titanium |  |  | titanium.html |
| 17 | Adhesive<br>resin cement | Duo-Link<br>Universal™ | BISCO | A-19620K | https://www.bisco.com/duo-link-universal-/ |

## Notes

### Competing Interest Statement

The authors have declared no competing interest.

